# Training induced improvements in knee extensor force accuracy are associated with reduced vastus lateralis motor unit firing variability

**DOI:** 10.1101/2022.02.10.479871

**Authors:** Isabel A Ely, Eleanor J Jones, Thomas B Inns, Síobhra Dooley, Sarah B J Miller, Daniel W Stashuk, Philip J Atherton, Bethan E Phillips, Mathew Piasecki

## Abstract

**Background:** Muscle force output during sustained submaximal isometric contractions fluctuates around an average value and is known to be influenced by variation in motor unit (MU) firing rates. MU firing rate variability seemingly reduces following exercise training interventions, however, much less is known with respect to peripheral MU properties. We therefore investigated whether targeted force accuracy training could lead to improved muscle functional capacity and control, in addition to determining any alterations of individual MU features.

**Methods:** Ten healthy participants (7 females, 3 males, 27±6 years, 170±8 cm, 69±16kg) underwent a 4-week supervised, unilateral, force accuracy training intervention. The coefficient of variation for force (FORCE^CoV^) and sinusoidal wave force tracking accuracy (FORCE^Sinu^) were determined at 25% maximal voluntary contraction (MVC) pre- and post-training. Intramuscular electromyography was utilised to record individual MU potentials from the vastus lateralis (VL) muscles at 25% MVC during sustained contractions, pre- and post-training.

**Results:** Knee extensor muscle strength remained unchanged following training, with no improvements in unilateral leg-balance. FORCE^CoV^ and FORCE^Sinu^ significantly improved in only the trained knee extensors by ~13% (*p*=0.01) and ~30% (*p*<0.0001) respectively. MU firing rate variability significantly reduced in the trained VL by ~16% (*n*=8; *p*=0.001), with no further alterations to MU firing rate or neuromuscular junction transmission instability.

**Conclusion:** Our results suggest muscle force control and tracking accuracy is a trainable characteristic in the knee extensors, which is likely explained by the reduction in MU firing rate variability apparent in the trained limb only.

## Background

The human motor unit (MU) is the final component of the neuromuscular system, comprising of a single somatic motor neuron, including its axon, distal axonal branches, neuromuscular junctions (NMJ), and associated innervated skeletal muscle fibres. The MU is fundamental to muscle force generation (Heckman & Enoka, 2012), and motor output is driven by supraspinal commands, spinal reflex pathways and neuromodulation (Khurram et al., 2021), with populations of MUs being recruited and discharging at differing firing times. Thus, modulation of MU firing rate contributes to the increase and decrease of muscle force.

During muscle contraction, the desired force output fluctuates around an average value rather than being at a constant level (Enoka et al., 2003; Enoka & Farina, 2021). Variation in MU firing rate has been identified as a critical determinant influencing the control of muscle force (Vila-Cha & Falla, 2016; Enoka & Farina, 2021), with associations between muscle force and MU firing rate dependent on single MU force, the input-output function of motor neurons and the frequency response of the muscle to transform an activation signal into force (Enoka & Farina, 2021). Using computational models allowing manipulation of key MU parameters (MU firing rate and MU firing rate variability), increasing index finger force resulted in MU firing rate variability of the flexor dorsal interosseous reducing exponentially, corresponding with improved simulated force fluctuations (Moritz et al., 2005). These simulated data are consistent with experimental (Laidlaw et al., 2000) and other simulated observations (Enoka et al., 2003), supporting evidence that MU firing rate variability (i.e., the variability of inter-discharge intervals across consecutive MU firings) is a, if not the, key physiological parameter influencing the ability to maintain steady muscle contractions (Vila-Cha & Falla, 2016). Compared to central MU function, much less is known with respect to peripheral MU features (i.e., NMJ transmission instability) and the influence these may have on muscle force control. Peripheral factors such as the release of acetylcholine at the NMJ, sodium/potassium pump activity, or modification to sodium and/or potassium intracellular and/or extracellular concentrations may alter muscle fibre action potential transmission (Allen et al., 2008). Although this alteration may subsequently impact muscle contraction and thus levels of force control, this has not yet been explored in a longitudinal manner.

The coefficient of variation for force (FORCE^CoV^) has been identified as a significant explanatory variable for multiple performance tasks including balance (Zech et al., 2010), walking (Davis et al., 2020a), manual dexterity (Kornatz et al., 2005; Keogh et al., 2019), levels of tremor (Kavanagh et al., 2016; Keogh et al., 2019) and the risk of falling in older adults (Carville et al., 2007; Enoka & Farina, 2021). The use of exercise training strategies (e.g., resistance exercise training (RET)) to improve muscle force control is, therefore, of interest for multiple diverse groups of individuals, including athletes, older adults, and those who are clinically vulnerable. Although RET is known to improve muscle strength, the impact on muscle force control is less equivocal. For example, Beck et al., reported that while improvements in knee extensor maximal voluntary contraction (MVC) force were observed following 8 weeks RET (~80% 1-repetition maximum (1RM)) in young individuals, neither FORCE^CoV^ or common drive were altered (Beck et al., 2011). Conversely, performing light-load (30% 1RM) training led to improvements in both knee extensor strength and knee extensor and elbow flexor muscle force control (Kobayashi et al., 2014), with the greatest RET-induced improvements in isometric FORCE^CoV^ occurring in the least steady subjects (Tracy & Enoka, 2006). Offering a potential explanation for improvements in FORCE^CoV^ with RET, 4 weeks of isometric strength training which significantly increased muscle strength also increased MU firing rate (+3±2.5pps) during the plateau phase of submaximal muscle contractions and decreased in the MU recruitment-threshold (Del Vecchio et al., 2019). Despite RET proving to be mostly an effective training mechanism to improve FORCE^CoV^, RET may not be accessible to all individuals due to physical limitations (Barry & Carson, 2004), meaning that alternative training modalities, with a focus on light-load/task specific training, still need to be established.

The aim of the current study was, therefore, to investigate the effect of a 4-week low intensity force accuracy training strategy on levels of knee extensor muscle force control/accuracy and any subsequent alterations to central and peripheral MU function in the vastus lateralis (VL) muscle. We hypothesised that muscle FORCE^CoV^ and sinusoidal wave tracking accuracy (FORCE^Sinu^), but not muscle strength, would improve with this training strategy, and that reduced MU firing rate variability would be observed in the trained VL.

## Methodology

### Participant characteristics

This study was approved by the University of Nottingham Faculty of Medicine and Health Sciences Research Ethics Committee (Reference number: C16122016) and was conducted in accordance with the Declaration of Helsinki. Written informed consent was obtained prior to participation in the study. Participants were excluded from the study if they presented with any musculoskeletal injury or neurological conditions. Ten healthy participants (7 females, 3 males, 27±6 years, 170±8 cm, 69±16 kg) recruited from the University of Nottingham staff and student population completed this study, each of whom were required to undergo 2 assessment visits (pre- and post-training) separated by 4 weeks of unilateral (right leg) force accuracy training. Participants arrived at the laboratory for assessment visits at ~0900 h (±1 h) having undergone an overnight fast (>10 h), with post-training testing occurring ~48 h after the last training session. All experimental techniques were completed bilaterally on assessment days.

### Assessment of muscle strength, FORCE^CoV^, FORCE^Sinu^ and unilateral balance

Knee extensor muscle strength was assessed via maximal voluntary contractions (MVC). Participants sat in a purpose-built dynamometer with hips and knees flexed at ~90° and each leg secured via a non-compliant strap to a force transducer just above the malleoli. After a standardised warm-up (10 sustained contractions held for 5-10 seconds, at ~50% MVC), participants were instructed to perform an MVC. Three attempts were made, each separated by 60 seconds, with real-time visual feedback (using Spike2 software v9.06) and strong verbal encouragement provided. The highest peak force (N) was determined as maximal.

FORCE^CoV^ was quantified from a set target line of 25% MVC (Figure 1A). Participants were given a single familiarisation trial before performing 6 contractions at this intensity. Each contraction lasted 12-15 seconds with a rest of 30 seconds between contractions. To avoid corrective actions when reaching the target line, the first two passes were excluded from the calculation. From these 6 contractions, the mean FORCE^CoV^ was subsequently calculated as ((standard deviation/mean)*100) from the plateau phase of the contraction.

**Figure 1.**
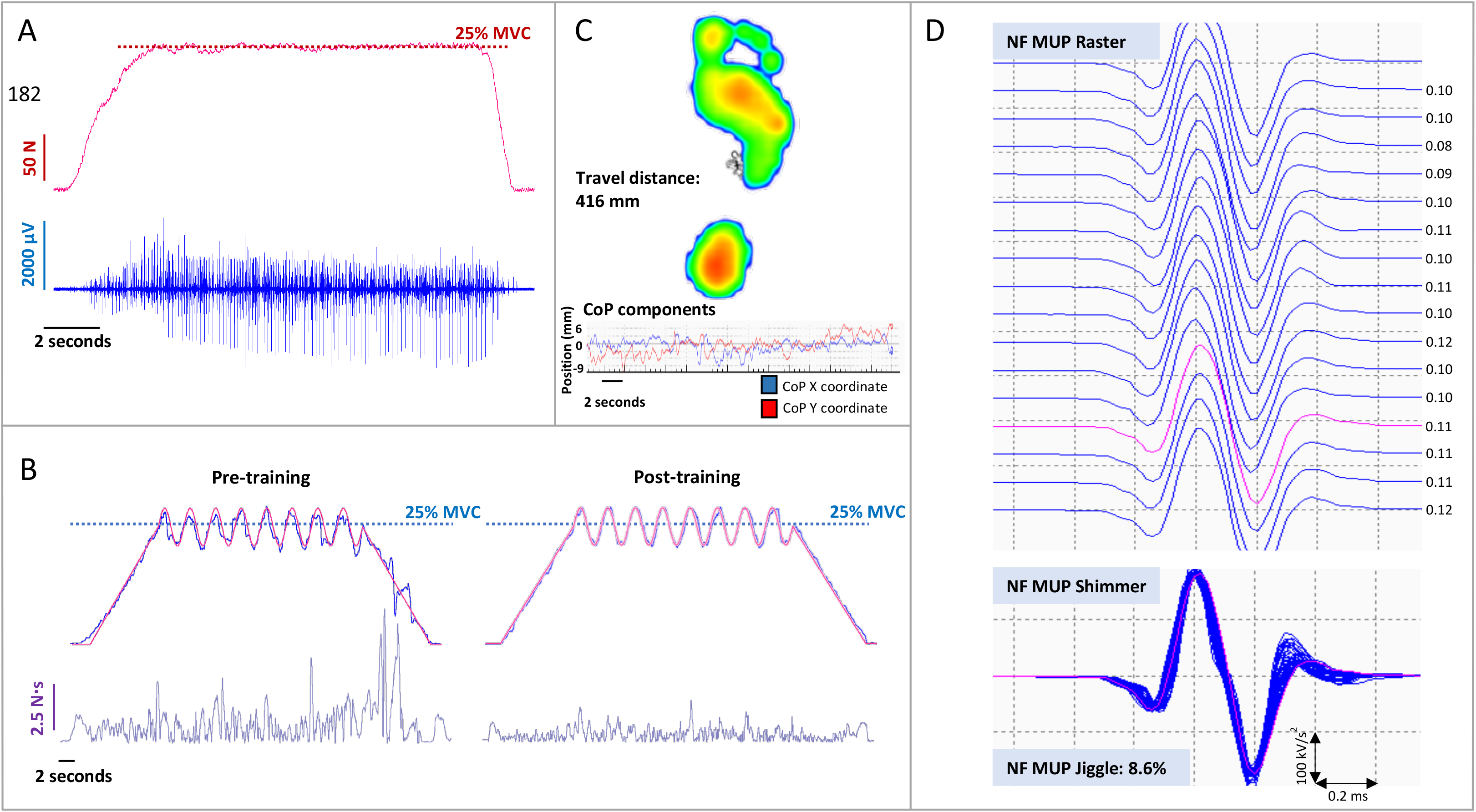
**(A)** Example force and intramuscular electromyography (iEMG) data recorded during a sustained isometric muscle contraction at 25% maximal voluntary contraction (MVC) in the vastus lateralis muscle; **(B)** Example raw data from a complex force tracking task at 25% MVC pre- and post-training. The red line represents the requested target force, whilst the blue line represents the observed force. Subsequent calculation of area under the curve (N.s) allowed for quantification of force tracking accuracy following training; **(C)** Unilateral balance data allowing measurement of centre of pressure (CoP), and the displacement of this (travel distance, mm), during static one-legged standing; **(D)** Example Raster and Shimmer plots of near fibre motor unit potentials (NF MUP) extracted from decomposed iEMG recordings from the vastus lateralis muscle during 25% MVC, allowing quantification of NF MUP jiggle; an indicator of neuromuscular junction transmission instability. Inter-discharge interval timings (ms) are indicated for each NF MUP firing in the motor unit potential train.

Participants next completed a series of sinusoidal wave force tracking tasks (using OT Bioelettronica software, Torino, Italy) at 25% MVC to assess levels of FORCE^Sinu^. A familiarisation contraction was performed prior to the assessment contraction. Contractions consisted of 8 oscillations at a set amplitude (±4%) lasting for 30 seconds. A 10-second ramp preceded and followed each oscillating section of the contraction to allow force to steadily increase and decrease to and from the desired contraction intensity (Figure 1B). Contractions were exported and analysed in Spike2 (Version 9) software, where a virtual channel was created (via subtracting the performed path from the requested path, and rectifying) and the area under the curve (AUC, N.s) of this channel was representative of the level of deviation from the target line, reflecting muscle force tracking accuracy.

Participants also completed physical function tests to assess unilateral balance of both legs pre- and post-training. All balance tests were performed using a Footscan plate (Footscan, 200 Hz, RSscan International, Belgium) allowing measurement of centre of pressure, and the displacement of this, during static one-legged standing. Participants were asked to visually focus on a fixed point in front of them for the duration of the test (30 seconds). A 5-second countdown was given before instruction to lift one leg, 2 seconds before the recording period began. Distance travelled (mm), the displacement of centre of pressure, was recorded for further analysis (Figure 1C).

## Intramuscular EMG measures

### Motor point identification

The motor point of each muscle was identified as the site of the muscle that produced the largest localised visible twitch using a low stimulation current with a cathode probe (Medserve, Daventry, UK) and a self-adhesive anode electrode (Dermatode, Fermadomo, BR Nuland, the Netherlands) (Piasecki et al., 2016a). A constant current stimulator (Digitimer DS7AH, Digitimer, Welwyn Garden City, Hertfordshire, UK) was set to a compliance voltage of 400 V with a 50 μs pulse width.

### Sampling of single motor units during voluntary contractions

Intramuscular electromyography (iEMG) recordings were obtained using disposable concentric needle electrodes with a recording area of 0.07 mm^2^ (Model N53153: Teca, Hawthorne, NY), with a grounding electrode on the patella. Participants were asked to relax their muscles to enable insertion of the needle electrode to enable sampling of MUs during the series of voluntary isometric contractions used to assess FORCE^CoV^ (as described above). Following each contraction, the needle electrode was withdrawn 5-10 mm and the bevel rotated 180°, recording from a total of 4-6 contractions from spatially distinct areas (Jones et al., 2021). iEMG signals were sampled at 50 KHz and bandpass filtered at 10 Hz to 10 KHz. Signals were digitised with a CED Micro 1401 data acquisition unit (Cambridge Electronic Design Ltd, Cambridge, UK). All iEMG and force signals were recorded and displayed in real-time via Spike2 software (CED, Cambridge, UK, version 9). Data were analysed offline in Spike2 (v9.06) software.

### iEMG signal analysis

iEMG data are available for 8 participants, as too few motor unit potentials (MUPs; <4) were isolated in the control limb post intervention in 2 participants. Procedures for identifying, recording, and analysing individual MUPs and calculating near fibre (NF) parameters have been described in detail elsewhere (Stashuk, 1999; Piasecki et al., 2016b). In brief, decomposition-based quantitative electromyography software was used to analyse the iEMG signal and visual inspection of all MUPs was performed. Sets of individual MUPs generated by a single MU were identified in the iEMG signal and extracted into MUP trains (MUPT). MUPTs composed of MUPs generated by more than one MU or having fewer than 40 MUPs were excluded.

Individual MUPs within a MUPT were further used to assess MU firing rate and NF jiggle. MU firing rate is expressed in Hz. MU firing rate variability was calculated as the coefficient of variation of the MU inter-discharge-interval times and expressed as a percentage. A NF MUP (Figure 1D) was obtained by calculating the slope of its corresponding MUP, using a low-pass second-order differentiator (Piasecki et al., 2021). This effectively reduces the uptake area of the needle electrode thus ensuring only near fibres significantly contribute to a detected NF MUP. NMJ transmission instability is quantified as NF jiggle, a measure of the variability of consecutive NF MUP shapes across a train (Stålberg & Sonoo, 1994, Piasecki et al., 2021). A total of 1,271 MUPs were recorded from all participants across all contractions from both legs and timepoints (pre- and post-training). This consisted of a total of 649 in the trained limb and 622 in the untrained limb. A mean of 8±3 MUPs were sampled with each needle location.

### Force accuracy training

All participants were required to perform force accuracy training 3x/week for 4 weeks, with all sessions being fully supervised by a member of the research team. Training was completed unilaterally, with all participants training the right leg which was also the dominant limb for all volunteers. Participants completed 6 sinusoidal force tracking contractions (as described previously and illustrated in Figure 1B) for each training session, with contractions conducted at 10% MVC (x2), 25% MVC (x2) and 40% MVC (x2), determined in the pre-training assessment visit, in a randomised order. The number of oscillations (either 6, 8 or 10) and amplitude (either ±2%, ±4% or ±8%) of the sinusoidal waves was different to that used in experimental testing (as stated above) and was again randomised for each training session. Contractions lasted 20-30 seconds dependent on contraction intensity and were separated by 60 seconds rest.

### Statistical analysis

Data are presented as mean ± SD unless stated otherwise. Muscle strength, FORCE^CoV^, FORCE^Sinu^ and distance travelled during unilateral leg balance were analysed using a two-way repeated measures ANOVA (condition*time) to assess measures in both the trained and untrained legs pre- and post-training. Bonferroni post hoc tests were used to identify statistical differences as a result of training. Multi-level mixed effects linear regression models were used to analyse MU data at 25% MVC with leg (trained *vs*. untrained) and time (pre *vs*. post) as factors. Interactions were first examined, and where present, individual coefficients for each leg are reported. Where no interactions existed, they were removed from the model and main effects of each leg were assessed individually. Adjusted beta values and 95% confidence intervals (CIs) are reported for all parameters. Statistical significance was accepted at *p*<0.05. Data analysis was performed using GraphPad Prism Version 8 (GraphPad Software, CA, USA) for analysis involving two-way repeated-measures ANOVA, and STATA SE Version 16 (StataCorp, Texas, USA) for multi-level mixed effects regression.

## Results

### Muscle strength and unilateral balance

Knee extensor maximal isometric strength remained unchanged following 4 weeks unilateral force accuracy training (Figure 2A; condition*time interaction effect: *p*=0.39) in both the trained (464.4±173.4 N *vs*. 447.0±173.1 N, *p*=0.97) and untrained (450.6±171.6 N *vs*. 403.0±160.7 N, *p*=0.13) legs.

**Figure 2.**
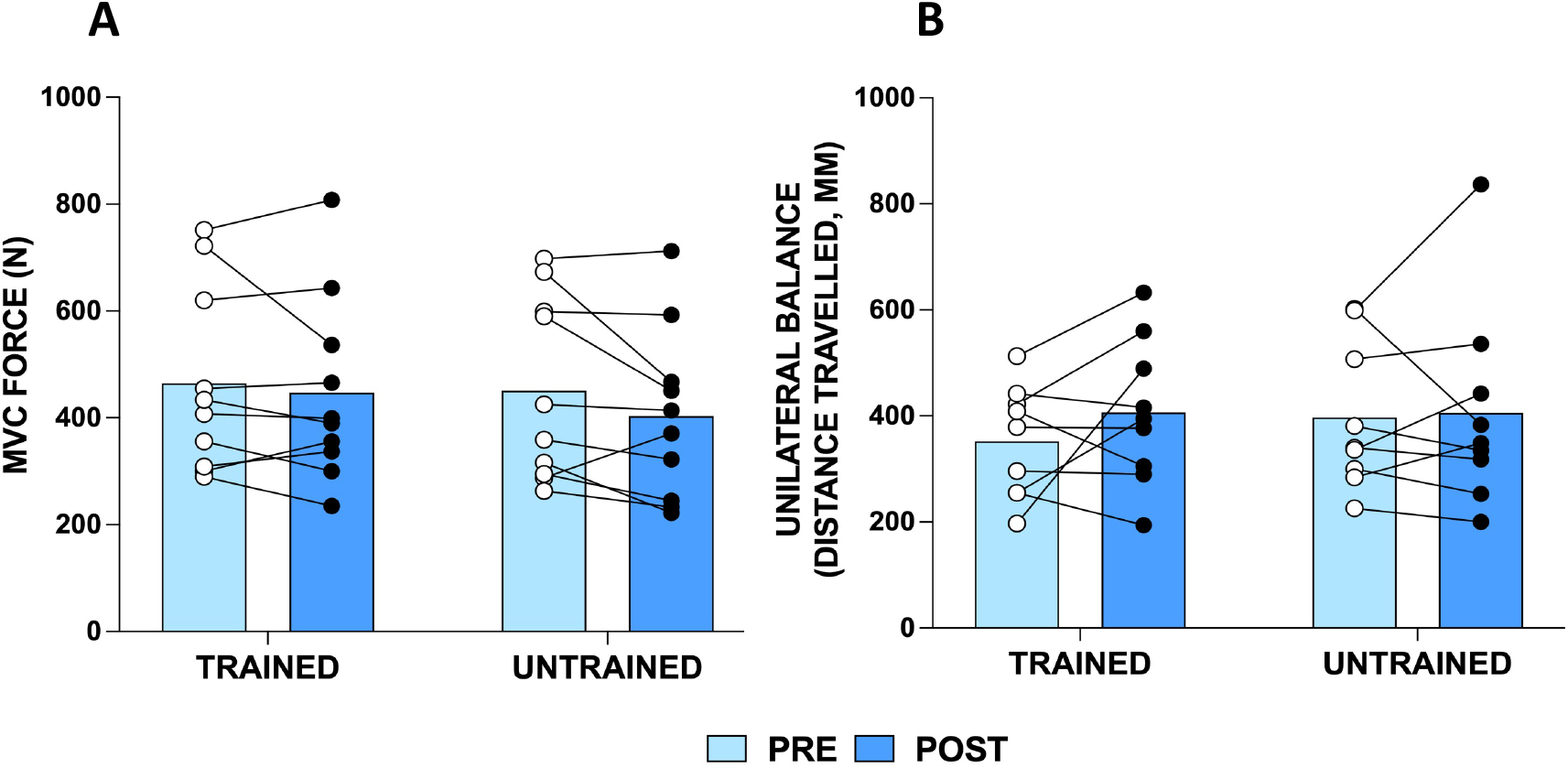
**(A)** Knee extensor maximum voluntary contraction (MVC) force (N), and **(B)** displacement of centre of pressure during unilateral balance tasks in n=10 young individuals (n=9 for unilateral balance due to one participant not achieving the minimum balance time required time to complete the test) pre- and post-training in the trained and untrained limb. Group mean shown as bars with individual data overlaid.

Displacement of centre of pressure during unilateral static standing did not change, thus no improvements in unilateral balance were observed, in either leg following force accuracy training (Figure 2B; n=9; trained: 352.0±105.4 mm *vs*. 406.6±137.8 mm, *p*=0.42; untrained: 397.2±138.6 mm *vs*. 405.9±189.2 mm, *p*>0.99).

### Muscle force tracking accuracy

Although the interaction effect was not significant (*p*=0.053), the trained knee extensors showed a significant improvement in FORCE^CoV^ following force accuracy training (trained: (2.80±0.58% *vs*. 2.39±0.40%, *p*=0.01; untrained: 3.02±0.71% *vs*. 3.00±0.65%, *p*>0.99; Figure 3A).

**Figure 3.**
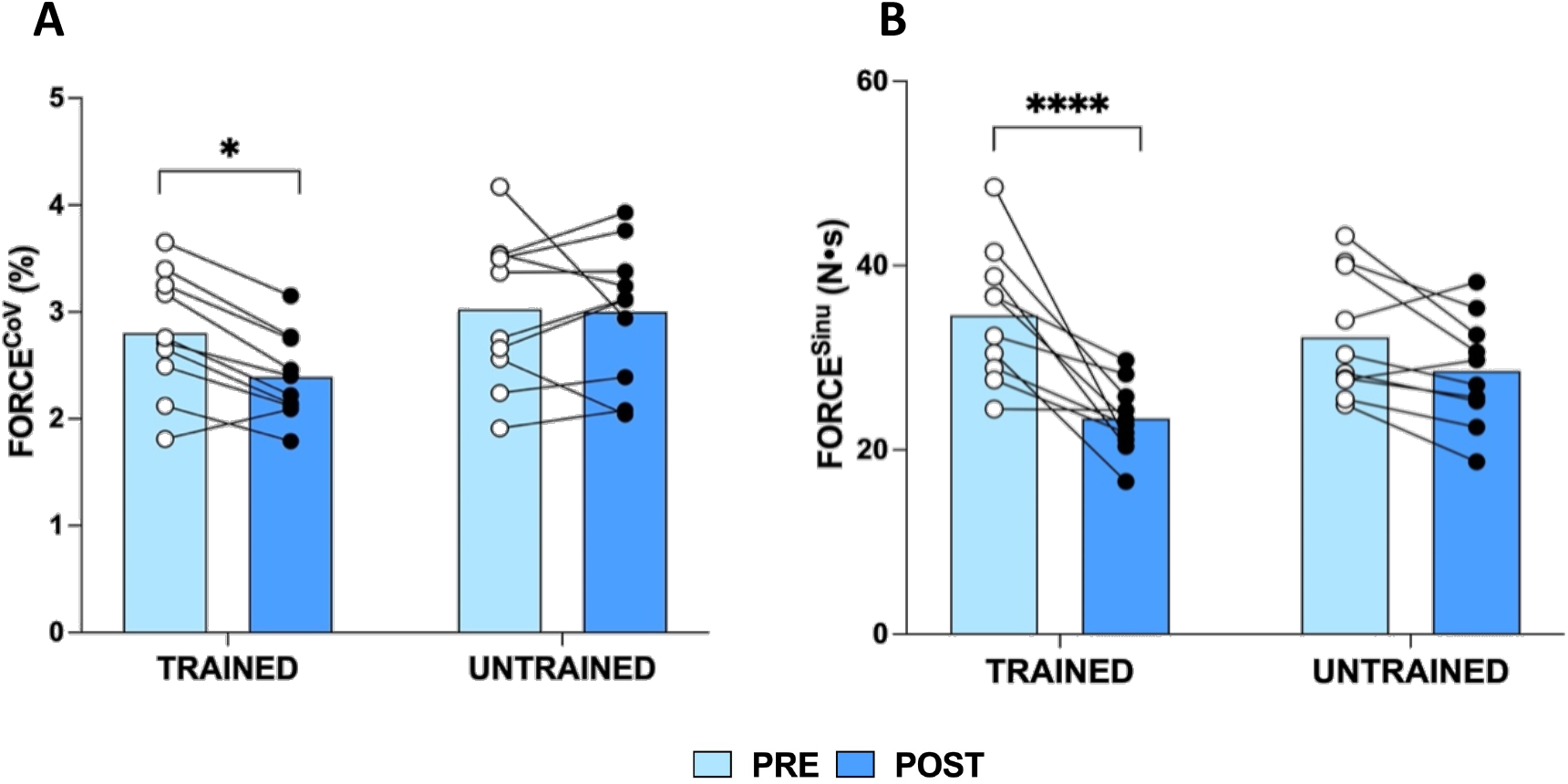
**(A)** Knee extensor coefficient of variation for force (FORCE^CoV^, %), and **(B)** knee extensor sinusoidal wave force tracking accuracy (FORCE^Sinu^; calculated as area under the curve, AUC (N.s)) at 25% maximal voluntary contraction (MVC) pre- and post-training in the trained and untrained legs of n=10 young individuals. Group mean shown as bars with individual data overlaid. *, *p*<0.05; ****, *p*<0.0001.

A significant interaction effect (condition*time, *p*=0.02) was observed for knee extensor FORCE^Sinu^ (Figure 3B), with improvements in FORCE^Sinu^ in the trained (34.59±7.26 N.s *vs*. 23.40±3.85 N.s, *p*<0.0001) but not the untrained (32.22c6.76 N.s *vs*. 28.57±5.93 N.s, *p*=0.19) leg following force accuracy training.

### Motor unit features

VL MU firing rate variability (Figure 4A) displayed a significant interaction effect (*p*=0.041) following training, with decreased MU firing rate variability in the trained leg (ß=−2.018, 95% CI=[−3.202]-[−0.835], *p*=0.001) but no change in the untrained leg (ß=0.862, 95% CI=[−0.271]-[1.995], *p*=0.14). No changes were observed in VL MU firing rate (Figure 4B; trained leg: ß=0.011, 95% CI=[−0.319]-[0.541], *p*=0.613; untrained leg: ß=0.009, 95% CI=[−0.383]-[0.401], p=0.97) or NMJ transmission instability in either leg following force accuracy training (Figure 4C; trained leg: ß=-0.714, 95% CI=[−1.823]-[0.399], *p*=0.208; untrained leg, ß=0.239, 95% CI=[−0.856]-[1.324], *p*=0.67).

**Figure 4.**
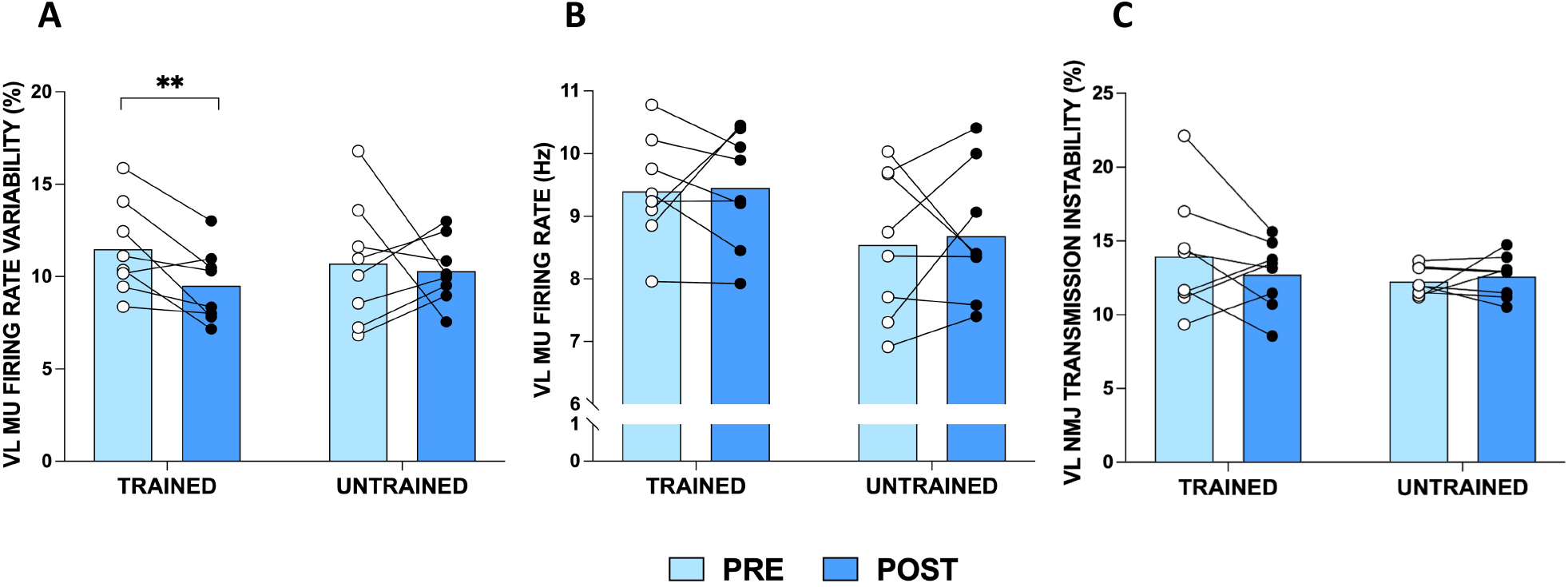
**(A)** Vastus lateralis (VL) motor unit (MU) firing rate variability (%), **(B)** VL MU firing rate (Hz), and **(C)** VL neuromuscular junction transmission (NMJ) instability (%) at 25% maximal voluntary contraction, pre- and post-training (*n*=8), in the trained and untrained legs. Group means shown as bars with individual data overlaid for data visualisation only. All analyses were based on multi-level linear regression models, where MUs were clustered to each muscle/participant. **, *p*=0.001.

## Discussion

The aim of the current study was to investigate the effects of a force accuracy training programme, with respect to muscle FORCE^CoV^ and alterations to individual MU features in VL. We highlight that 4 weeks of force accuracy training resulted in improvements in muscle force accuracy and control, evidenced through improved FORCE^CoV^ and FORCE^Sinu^ in the trained knee extensors only. No changes in muscle strength or displacement of pressure during unilateral balance tasks were observed with this form of training. The improvements in force accuracy and control were associated with decreased MU firing rate variability, again occurring in the trained limb only.

In line with our hypothesis, MVC force remained unchanged in the knee extensors across both legs. However, these results are not in direct agreement with some others. For example, in older adults, low intensity RET (30% 1RM) increased knee extensor MVC after both 8 (Kobayashi et al., 2014) and 16 (Tracy & Enoka, 2006) weeks of training, with numerous studies reporting that low intensity RET (i.e., ~20-40% 1RM) can induce increases in muscle strength (Keen et al., 1994; Hortobagyi et al., 2001; Tracy & Enoka, 2006; Kobayashi et al., 2014). This notion does however remain inconclusive with studies such as that by Moore et al., reporting unchanged isometric elbow flexor strength following 8 weeks RET in young individuals at 50% 1RM (Moore et al., 2004). Considering the current study, we suggest an absence of increased strength following our force accuracy training may have occurred due to: i) a fixed training load (i.e., %MVC was not increased throughout training), and ii) studying a population of healthy young individuals, with previous studies of similar methodology often recruiting previously sedentary older adults; a population arguably having the most to gain from being physically active (McPhee et al., 2016).

Although we demonstrate no functional improvements in the displacement of centre of pressure during unilateral balance tasks post-training, FORCE^CoV^ has previously been correlated with balance measures (Kouzaki & Shinohara, 2010; Oshita & Yano, 2010; Davis et al., 2020b). For example, FORCE^CoV^ assessed via low intensity isometric contractions in the plantar flexors was significantly associated with balance performance with eyes closed (determined by the time, in seconds, to complete the test) in both young and older individuals. However, one study demonstrated this association for contractions at 20%, but not 10% MVC (Oshita & Yano, 2010), and another for contraction intensities ≤5% MVC (determined via displacement of centre of pressure) (Kouzaki & Shinohara, 2010). Davis and colleagues also demonstrate FORCE^CoV^ of the hip abductors and dorsiflexors to be the most significant explanatory variable during light load contractions to explain variance in sway-area rate across most balance conditions, although most of the variance across conditions was unexplained; suggesting other physiological mechanisms important for postural control likely influence unilateral balance (Davis et al., 2020b). The effects of force accuracy training on functional outcomes such as unilateral balance remains, therefore, to be further examined in other muscle groups and populations such as older individuals, who display deterioration of unilateral balance with advancing age (Maki et al., 1990; Izquierdo et al., 1999; Hess & Woollacott, 2005).

The current study utilised two different force tracking tasks, varying in difficulty, to assess levels of FORCE^CoV^ and force tracking accuracy. Our results demonstrate improvements in both FORCE^CoV^ and FORCE^Sinu^ following training. During isometric contractions of fluctuating force (i.e., sinusoidal contractions), the recruitment and subsequent de-recruitment of Mus needs to be aligned to match the desired trajectory to increase and decrease force (Duchateau & Enoka, 2008). As such, both MU recruitment and de-recruitment requires different control strategies (Duchateau & Enoka, 2008). Resultingly, the fluctuation in force during the sinusoidal contractions may contribute to positive alterations of force tracking and control strategies, hence observing improvements following training.

Commonly, FORCE^CoV^ has been highlighted as a critical explanatory variable with respect to muscular performance of tasks including walking (Davis et al., 2020a), tremor (Kavanagh et al., 2016; Keogh et al., 2019), and risk of falls in older adults (Carville et al., 2007), with FORCE^CoV^ progressively deteriorating from middle to older age (Piasecki et al., 2021). A recent study by Davis and colleagues examined plantar-flexor and dorsiflexor FORCE^CoV^ at 10 and 20% MVC in individuals who presented with multiple sclerosis and those who did not (Davis et al., 2020a). Patients with multiple sclerosis presented with worse FORCE^CoV^ in the dorsiflexors at 20% MVC, in addition to a reduced walking distance during a 6-minute walk test. In addition, in the patient group an increased time to walk 25 feet was associated with a greater FORCE^CoV^ during submaximal contractions in both muscles (Davis et al., 2020a). Importantly, a reduced walking distance in the control group was also associated with dorsiflexor FORCE^CoV^ and variation of common input to the plantar flexors during steady contractions (Davis et al., 2020a). Although multiple sclerosis is a neurological disorder impacting both the central and peripheral nervous systems (Davis et al., 2020a), the age-related decline in the neuromuscular system presents with aligned mobility-impairing symptoms which ultimately impact quality of life (Mitchell et al., 2012; Wilkinson et al., 2018). Therefore, the impact of force accuracy training on FORCE^CoV^ should be examined in older/clinically vulnerable populations to determine whether the more targeted approach (*vs*. traditional RET, for example) leads to improvements in force accuracy and control which may aid with performance of functional tasks.

No alterations to MU firing rate were noted in the present study which may be due to the low intensity of training as 4 weeks of strength training at ~75% MVC increased tibialis anterior MU firing rate during the plateau phase of submaximal isometric contractions (at 35, 50 and 70% MVC) and reduced MU recruitment thresholds (Del Vecchio et al., 2019). Similar effects on MU firing rate were also reported in the abductor digiti minimi when training was performed at a maximal intensity (Patten et al., 2001). Conversely, another study demonstrated increased VL MU firing rate at 100% MVC but not during submaximal contractions following 6 weeks of training using dynamic muscle contractions (Kamen & Knight, 2004), whilst 2 weeks of force modulation training (i.e., neuromuscular control training) reduced MU firing rate during contractions at 30-60% MVC (Patten & Kamen, 2000). Taken together, it is likely that adaptation to MU firing rate is muscle and contraction level specific, and sensitive to the form of training intervention.

In line with our hypothesis and the work of others (Laidlaw et al., 2000; Enoka et al., 2003; Moritz et al., 2005; Enoka & Farina, 2021), although not unanimous (Beck et al., 2011), we demonstrate MU firing rate variability to significantly reduce following force accuracy training. In flexor dorsal interosseous, Kornatz and colleagues highlight a weak association between reduced force fluctuations, in addition to improvements in manual dexterity, and declines MU firing rate variability (Kornatz et al., 2005), which was later strengthened using computational models (Moritz et al., 2005). Furthermore, MU firing rate variability was reported to reduce, as such enhancing knee extensor FORCE^CoV^, following strength but not endurance training; suggesting different training modalities result in varying neuromuscular adaptations (Vila-Cha & Falla, 2016). MU firing rate variability and force fluctuations are governed by central descending pathways modulated by independent and common synaptic inputs to the MU pool (Taylor et al., 2003; Vila-Cha & Falla, 2016), and here we demonstrate improvements in force accuracy and MU firing rate variability in the trained limb only, which may be a result of reduced antagonist muscle activity and/or inhibitory afferent feedback (Enoka & Farina, 2021). There is potential for ionic changes (e.g., modification to sodium and/or potassium ion intracellular and/or extracellular concentrations, subsequently altering muscle fibre action potential transmission (Allen et al., 2008)), release of acetylcholine at the NMJ, and/or the type/intensity of muscle contraction (Enoka et al., 2003; Carville et al., 2007; Enoka & Farina, 2021) to also influence levels of muscle force control, however we evidenced no alterations to NMJ transmission instability, assessed via NF MUP jiggle.

### Strengths and limitations

As the training period was only 4 weeks, it offers translational relevance and application to pre/rehabilitation scenarios (Durrand et al., 2019), and the short duration of each training session (~20 minutes) counters one of the most commonly cited barriers to exercise interventions; “lack of time” (Trost et al., 2002). Secondly, the tasks constituting force accuracy training are arguably more applicable to daily movements (e.g., rising from a chair) than traditional RET due to the fluctuations in force requiring greater muscle coordination. The use of iEMG and near fibre analysis to identify single MU features from a range of muscle depths in both male and female participants affords greater insight into central and peripheral adaptions at a single MU level than commonly achieved with surface electromyography methodology. A limitation of the study is that all MUs were sampled at 25% MVC which reveals little of adaptations that may be occurring with higher threshold MUs. Furthermore, despite providing great detail, an inherent limitation of iEMG is the inability to reliably track the same MUs pre- and post-intervention (Martinez-Valdes et al., 2017). Finally, reduced antagonist co-activation cannot be ruled out in the observed improvements in force tracking accuracy (De Luca & Mambrito, 1987).

## Conclusion

To summarise, we highlight that a 4-week period of targeted force accuracy leads to improved muscle force control and accuracy in young healthy participants, which is associated with reduced MU firing rate variability, as assessed by the coefficient of variation for the interspike interval. Importantly, these adaptations and possible mechanisms were evident in the trained limb only. These findings may influence interventional strategies to improve force accuracy, including in older and clinical populations where such improvements may help with independence maintenance via improved performance of activities of daily living.

## Declarations Acknowledgements

The authors would like to thank Mrs Amanda Gates and Dr Paula Scaife for their assistance with the study. The authors would also like to thank the participants for their time and efforts in the study.

## Data availability statement

The datasets generated and analysed during the current study are available from the corresponding author upon reasonable request.

## Author Contributions

All authors contributed to the conception and design of the work. IAE, EJJ, TBI, SD and SBJM acquired the data and IAE analysed the data. IAE, DWS, PJA, BEP and MP drafted the manuscript and prepared the figures. All authors approved the final version of the manuscript. All authors agree to be accountable for all aspects of the work in ensuring that questions related to the accuracy or integrity of any part of the work are appropriately investigated and resolved. All persons designated as authors qualify for authorship, and all those who qualify for authorship are listed.

## Abbreviations

FORCE^CoV^: coefficient of variation for force
FORCE^Sinu^: sinusoidal wave force tracking accuracy
iEMG: intramuscular electromyography
MU: motor unit
MUP: motor unit potential
MUPT: motor unit potential train
MVC: maximal voluntary contraction
NF: near fibre
NMJ: neuromuscular junction
RET: resistance exercise training
VL: vastus lateralis
1RM: 1-repetition maximum.

